# nVenn2: faster, simpler generalized quasi-proportional Venn diagrams

**DOI:** 10.64898/2026.01.19.700279

**Authors:** Samuel Pis-Vigil, María González-Pereira, Magda R. Hamczyk, Víctor Quesada

**Affiliations:** Departamento de Bioquímica y Biología Molecular, Universidad de Oviedo-IUOPA, Oviedo, Spain; Departamento de Bioquímica y Biología Molecular, Universidad de Oviedo, Oviedo, Spain; Aarhus Institute of Advanced Studies, Aarhus University, Aarhus, Denmark; CIBERONC, Madrid, Spain

## Abstract

Proportional Venn diagrams provide a compact representation of the relationships between sets. Each relationship is represented with a region whose area reflects the number of elements shared by a given combination of sets. This means that the number of regions grows exponentially with the number of sets, which is why proportional Venn diagrams with more than five sets are cumbersome to interpret and seldom used. However, Venn diagrams with a large number of sets may still be legible if enough regions are empty and do not need be represented. Here, we present nVenn2, the second version of the nVenn algorithm, to create quasi-proportional Venn diagrams. This new version uses a different, more flexible approach which includes steps to minimize the complexity of the diagram. Thus, computation time for nVenn2 mainly grows with the number of non-empty diagram regions, rather than with the number of sets. This property allows users to create interpretable quasi-proportional Venn diagrams with large numbers of sets. The nVenn2 algorithm is freely available as an executable program, as a web page, as an R package (nVennR2) and as a Python package (nVennPy). All interfaces allow users to edit the appearance of the resulting diagram.

## 1 Introduction

Venn diagrams were first described in the nineteenth century as tools for logical reasoning [1]. John Venn started with ideas from Leonhard Euler and others, in combination with the logical system developed by George Boole, to represent complex propositions in a single figure. Venn himself noticed that “[b]eyond five terms it hardly seems as if diagrams offered much substantial help”, since the number of propositions doubles with each new term. It is interesting to notice that this original description is focused on how negative propositions can be represented by eliminating regions in the general diagram. As the original aim of this representation was to exhaust the logical propositions from a number of terms, eliminating regions in a diagram was never considered as a strategy to make it more readable.

As a trivial extension of their use in the field of logic, Venn diagrams have also been used to illustrate basic concepts of set theory, such as “union” and “intersection” [2]. This use emphasizes the contents of sets as a measure of how similar or dissimilar those sets are. In this context, it makes more sense to eliminate regions corresponding to inexistent relationships between sets. For instance, if two sets have no elements in common, we can erase their intersection and depict them as two disjoint circles.

Moreover, a Venn diagram can be drawn in such a way that the area of each region is proportional to the number of elements in that region. This representation provides a quick overview of the relationships between sets, expressed as the proportion of elements they share. In bioinformatics, obtaining this overview is a recurrent task. A typical use involves sets of genes that are identified in different conditions of a given experiment. The proportional Venn diagram of those sets allows researchers to quickly detect abnormal regions (e.g., “most of the genes identified in condition A are also identified in condition B, but not in C”) and to obtain a list of the elements in each region (e.g. “which genes were identified in conditions A and B, but not in C?”).

Representing proportional Venn diagrams with more than two sets is challenging, and cannot be done in the general case using convex shapes. However, multiple programs attempt to solve this task in some limited form [3], consistent with the importance of these diagrams in bioinformatics and other fields. Some of these programs, like eulerAPE [4], use regular shapes and limit the number of sets that may be represented. Others, like eulerr [5] and Edeap [6], use ellipses for sets and relax the proportionality constraint by using penalty functions. Other representations focus on optimizing and simplifying the topological properties of the diagram [7, 8].

Our own solution for proportional Venn diagrams, called nVenn, uses irregular shapes, which allow for more accurate diagrams without limiting the number of sets [9]. Thus, the first nVenn algorithm started with a general Venn diagram. Each region of the diagram contained a circle with an area that was proportional to the number of elements in that region. This starting representation was then simplified using a simulation that maintained its topological properties while adjusting the set lines to the circles. While the resulting figures were often useful, several limitations existed that were intrinsic to the algorithm.

First, the starting diagram was always identical, and therefore the final figure was always the same for a given input. Since some representations could be sub-optimal (i.e., unnecessarily complex), some diagrams could not be improved without changing the order of the sets, and even in that case an optimal figure might not be obtained. Second, the starting diagram did not take advantage of empty regions, which made the simulation time scale non-linearly with the number of sets. This meant that nVenn diagrams with more than seven sets were often impractical. While in most cases such a diagram would not be useful, there are exceptions, as the interpretability of a proportional Venn diagram depends on the number of non-empty regions rather than on the number of sets.

Here, we describe the second version of the nVenn algorithm, which features a heavily modified procedure to address previous limitations. Thus, nVenn2 gives a different result, with minimized local complexity, each time it is run.

## 2 Implementation

Like the previous version, nVenn2 is coded in C++. To simplify the process of connecting different interfaces, new functions have been added to the core C++ libraries. The input of nVenn2 is a table where each row or column, as decided by the user, provides a set name and set elements. The corresponding description of the Venn diagram is then computed. In addition to running the simulation to get the diagram, the core libraries also allow users to explore the diagram by listing the elements corresponding to each region. An additional set of new functions can be used to edit the resulting diagram by changing colors, font sizes and other graphical features.

### 2.1 Steps

The final diagram is generated in seven steps (**Figure 1**). Each of these steps is finished when specific criteria have been met. This ensures that the simulation is not run for more steps than necessary.

**Figure 1:**
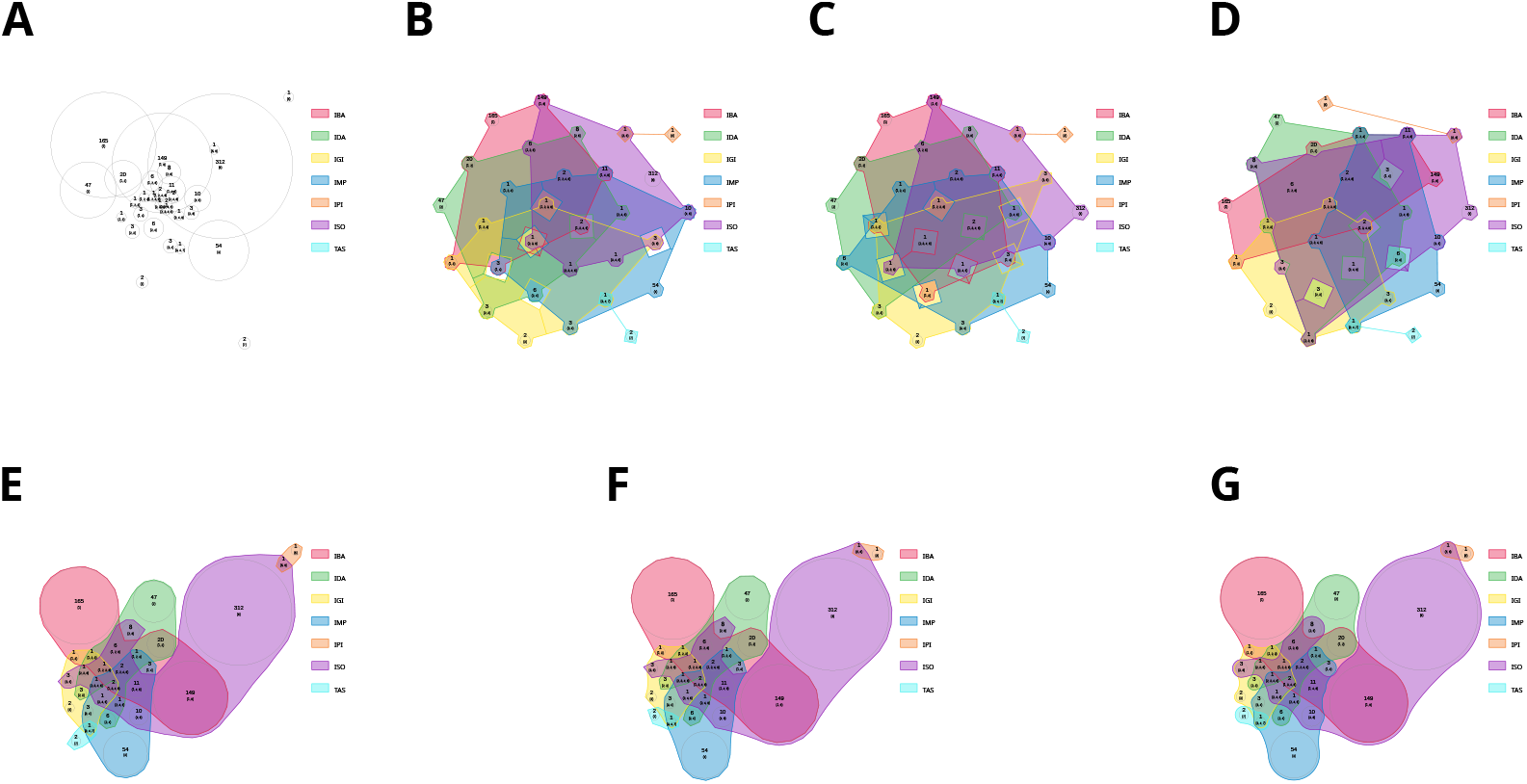
nVenn2 algorithm. **A**, circles representing regions are placed on grid pseudorandomly and find an equilibrium where similar regions are attracted and dissimilar regions are repulsed. **B**, a repulsive force separates circles into a new grid and a topologically correct Venn diagram is generated with lines surrounding all regions. **C**, regions are swapped pairwise to maximize a measure of how close similar regions are. **D**, regions are swapped pairwise to minimize the number of line crossings in the corresponding Venn diagram. **E**, the lines in the Venn diagram contract until regions are in contact. **F**, circles are packed closer together with attractive forces. **G**, circles are fixed in place and lines are smoothed.

In the first step (**Figure 1A**), each region is represented as a circle with an area that is proportional to the number of elements inside and placed pseudorandomly on a grid. Then, a simulation is run with a dissipative friction force and a spring-like force between each pair of circles. This spring-like force is stronger the more sets the corresponding regions have in common, and it becomes repulsive if the corresponding regions do not both belong to any common set. For instance, a circle (i.e., region) that belongs to sets 1, 2 and 4 will be attracted to the circle that belongs to sets 1 and 4 more strongly than to the region that belongs only to set 1, while it will be repulsed by the circle that belongs only to set 3. Once the total velocity of all circles is lower than a certain threshold, the second step sets a repulsive force between all the circles. The simulation is run until the minimum distance between circles is higher than a preset threshold (**Figure 1B**).

The two following steps minimize some property of the resulting grid. First, the value of the property is calculated. Then, the coordinates of two circles are swapped and the property is calculated again. If the new value is higher than the previous value, the swap is undone. Each step is repeated for every pair of regions until a certain number of steps has not improved the value of the main property (**Figure 1C,D**).

Once the minimization steps are completed, three additional steps simulate a physical system to reach a compact diagram (**Figure 1E-G**). These steps are similar to the simulation steps in the first version of nVenn [9] with some improvements in speed and stability. The most important change is that after each step a diagram property (compactness or area) is calculated and compared to that of previous steps. If this property is not consistently improving, the step is finished. More details about each step can be found in **Supplementary Data**.

### 2.2 Interfaces

In addition to direct compilation, nVenn2 can be used from a web page, R and Python. Each of these interfaces uses the C++ files without modifications, so that improvements to the libraries can be easily propagated.

#### 2.2.1 Web page

The web page uses HTML, CSS, vanilla javascript and SVG for the interface. The library itself was compiled into WebAssembly [10] using Emscripten v3.1.69 (https://emscripten.org/). The text table describing each set can be directly pasted into a text area or read from a file.

Once the diagram is generated user can explore it using tick boxes or directly clicking on regions. A text box will show the elements exclusively belonging to the provided region. A new version of the diagram can be created by simply clicking a button.

The interface contains several controls to rotate and modify the resulting figures. A diagram can be saved as an SVG file or standalone web page, or exported as a PNG (bitmap) file. Saved SVG or web page files can be reloaded into the interface for further processing. A tutorial provides step-by-step instructions to create, explore, edit and store diagrams.

#### 2.2.2 R library

The R interface uses Rcpp [11] to create function bindings to the library methods. Since a C++ object is not trivial to store in R, only equivalent R objects are saved. When those R objects are passed to Rcpp methods, the C++ library can read the code and internally rebuild the original C++ object. In keeping with the functional paradigm of R, all methods return a modified R object without changing the original object.

An advantage of Rcpp is that R types can be easily passed to C++ code. Therefore, this interface allows users to create an object from a text table and also from a list of lists, similar to the first version of nVennR. A vignette is provided with examples of use.

#### 2.2.3 Python library

The Python library contains bindings created with Pybind11 (https://github.com/pybind/pybind11). This interface uses the bindings directly to create a Python object that can be accessed and modified. The diagram contained in the object can be edited, saved and exported. This version only creates objects from text tables.

An interactive, publicly available Jupyter Notebook in Binder documents the functions with examples (https://mybinder.org/v2/gh/vqf/codespaces-jupyter/HEAD?urlpath=%2Fdoc%2Ftree%2Fnotebooks%2Fdoc.ipynb).

## 3 Testing

To compare the first and second versions of nVenn, we generated pseudorandom Venn diagrams in R and represented them with nVennR and nVennR2. Briefly, for each test a universe size was chosen. Then, each set was constructed by picking a pseudorandom number of elements from that universe. Since nVennR does not have a mechanism for ending execution, we used as many 5000-cycle execution blocks as necessary to ensure consistently compact diagrams. Testing code is available at https://github.com/uo283276/Pruebas_nVenn.

As expected, the performance of both algorithms was very similar with diagrams up to five sets (**Figure 2A**). For six or more sets, nVennR2 was consistently faster than nVennR. In fact, Venn diagrams with more than nine sets were feasible with nVennR2 and not with nVennR. We also represented the computational time needed relative to the number of regions in the diagram (**Figure 2B**). Again, execution times were similar for diagrams with few regions, although nVennR2 showed more consistency between runs. For higher numbers of regions, nVennR2 was clearly faster and much more consistent.

**Figure 2:**
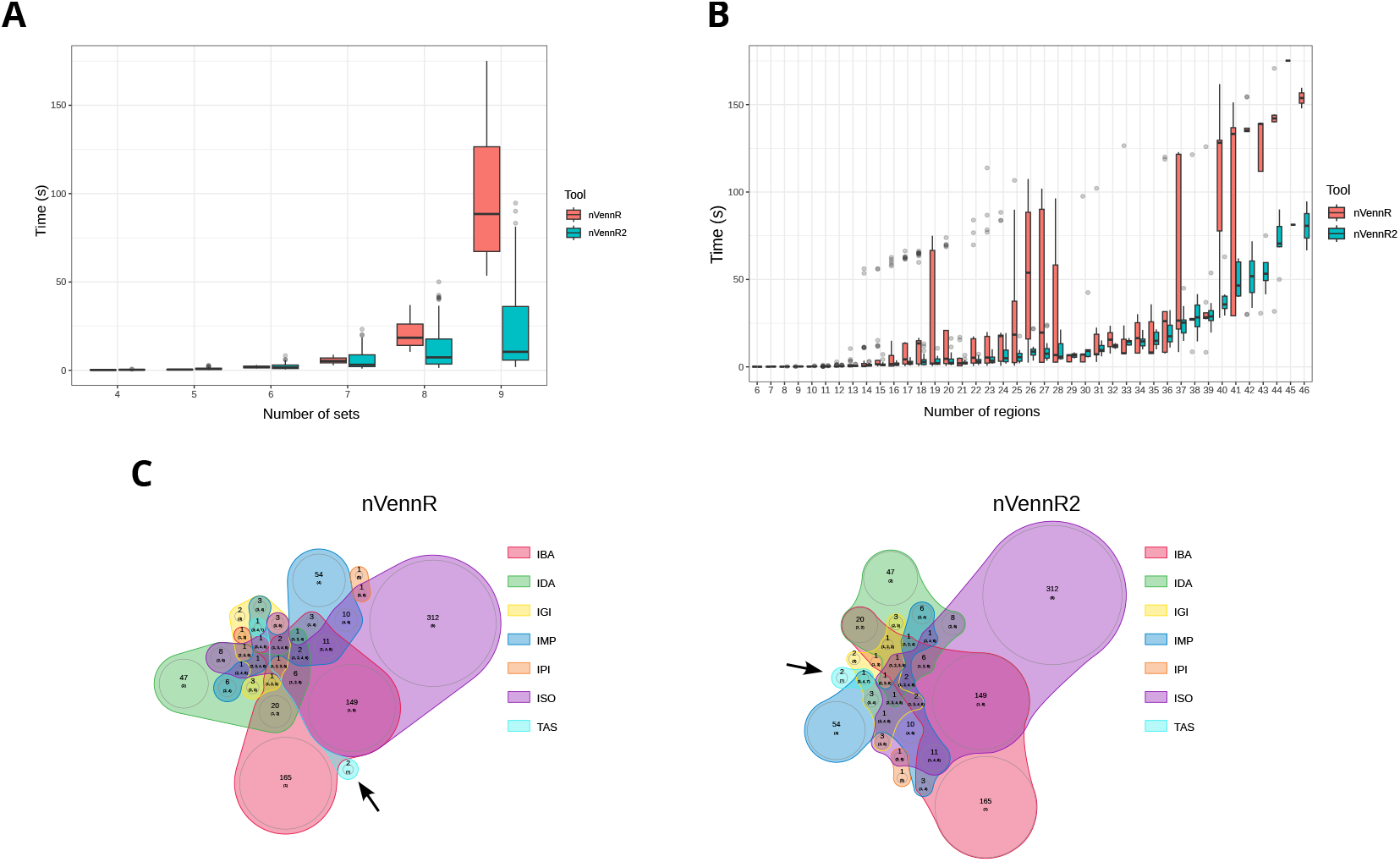
Performance of nVenn algorithms. **A**, Execution time for diagram completion with different numbers of sets. Boxes show the span from the first and third quartiles of the results, black horizontal lines show the median and whiskers represent an approximate 95 % confidence interval (N=40 for each number of sets). **B**, Execution time for diagram completion with different numbers of regions. Boxes show the span from the first and third quartiles of the results, black horizontal lines show the median and whiskers represent an approximate 95 % confidence interval (N=40 for each number of regions). **C**, seven-set diagram constructed with nVennR (*left*) and nVennR2 (*right*). Genes included in the innate immune system GO category (GO: 0045087). Seven subsets with different GO evidence codes were generated: IBA (Inferred from Biological aspect of Ancestor), IDA (Inferred from Direct Assay), IGI (Inferred from Genetic Interaction), IMP (Inferred from Mutant Phenotype), IPI (Inferred from Physical Interaction), ISO (Inferred from Sequence Orthology) and TAS (Traceable Author Statement). Arrows point to set 7 (TAS).

In addition, we found multiple examples of nVennR diagrams that were improved by nVennR2. For instance, the diagram in **Figure 1** performed with nVennR shows non-optimal placement of some regions (**Figure 2C**). Thus, the relationship of set seven (TAS) with the rest of sets is much easier to interpret with nVennR2.

## 4 Discussion

The aim of nVenn2 is to provide a faithful and aesthetically pleasing representation of the relationships between sets, as well as a set of tools to explore the corresponding regions. The representation itself follows the same criteria as the first version of the algorithm. Notably, this means that each region is represented by the corresponding circle, whose area is proportional to the size of the region. The set line surrounding those circles does not necessarily follow that proportionality. For this reason, nVenn2 does not provide any mechanism to alter the circles representing regions. Moreover, the radius of any circle may not be less than 2 percent of the width of the diagram, which means that the area of smaller regions may not faithfully represent the number of elements it contains.

The second version of nVenn features a complete rework of the original algorithm. Only the last three steps of nVenn2 have counterparts in the first version, and they have been streamlined. At the core of the new algorithm, a procedure generates a topologically correct diagram from a grid of circles representing regions. In particularly complex cases, this step can fail. However, we found no errors during testing with hundreds of Venn diagrams, which suggests that the error frequency is exceedingly low.

Another fundamental difference between versions is that nVenn2 yields a different diagram after every run. Users are encouraged to try several simulations to get a figure that best represents the data. With this feature, we have been able to balance exhaustiveness and running time. Thus, the topology of the diagram is decided using a form of gradient descent. While this is usually good enough for simple diagrams, in more complex situations the algorithm will likely stop at a local minimum. A procedure that looks for global minima in arbitrary cases would require large computational resources and running time.

It is also worth noting that the source of randomness in the code is the mt19937 Mersenne twister engine fed by the C++ standard random device. The definition of this random device allows implementations that yield the same number sequence every time. If the code is compiled with this implementation, the same input will yield the same output every time the program is started. In this case, repeated execution without closing the program will still produce different diagrams. The compilers for the provided interfaces do not have this limitation.

In summary, the second version of nVenn overcomes some of the most obvious limitations in the first version. In most cases, nVenn2 will be faster and provide more informative diagrams. In addition, we are providing multiple interfaces to allow for fast integration of nVenn2 with existing workflows.

## 5 Availability

All libraries and interfaces are publicly available under MIT licenses:

**nVenn2** https://github.com/vqf/nVenn

**Web interface** https://vqf.github.io/nvenn2/nvenn2.html and https://degradome.uniovi.es/vqf/nvenn2/nvenn2.html

**Python interface** https://github.com/vqf/nVennPy. Also available from Pypi (https://pypi.org/project/nvenn2/ or from the command line with “pip install nvenn2”).

**R interface** https://github.com/vqf/nVennR2

## 6 Acknowledgments

This work was funded by the Spanish Ministerio de Ciencia, Innovación y Universidades (PID2022-141488OB-I00). We thank Dr. David Roiz-Valle for his useful advice during testing.

