## Supplementary Methods for "nVenn2: faster, simpler generalized quasi-proportional Venn diagrams"

### Supplementary Data

Pis-Vigil *et al.*

January 12, 2026

#### Supplementary Methods

##### Tangent class

Several methods use a class called *tangent* to store a slope without ambiguities. The object is created with two values, representing a $\Delta x$ and a $\Delta y$ or, equivalently, two points $p_1$ and $p_2$, where $\Delta x = x_{p_2} - x_{p_1}$ and $\Delta y = y_{p_2} - y_{p_1}$. Both values are stored, as well as a *quadrant* and a *slope* calculated from those values:

If $(\Delta x < 0 \wedge \Delta y \leq 0)$: $Quadrant = 1$; $Slope = \frac{\Delta y}{\Delta x}$

If $(\Delta x \geq 0 \wedge \Delta y < 0)$: $Quadrant = 2$; $Slope = -\frac{\Delta x}{\Delta y}$

If $(\Delta x > 0 \wedge \Delta y \geq 0)$: $Quadrant = 3$; $Slope = \frac{\Delta y}{\Delta x}$

If $(\Delta x \leq 0 \wedge \Delta y > 0)$: $Quadrant = 4$; $Slope = -\frac{\Delta x}{\Delta y}$

This ensures that each angle has a unique quadrant/slope combination and that every value is finite. In addition, angles do not need to be computed with expensive arctan operations. Another useful feature is that this class establishes a quick comparison between objects, where higher values mean *more counter-clockwise* starting at the negative x axis. Thus, an object with a higher quadrant is higher than an object with a lower quadrant. If two objects belong to the same quadrant, the object with a larger slope is higher.

Another useful operation is subtraction. Let $t_1$ and $t_2$ be two tangent objects. The operation $t_2 - t_1$ returns a tangent object equivalent to $t_1$ after rotating both objects so that $t_2$ sits at quadrant 1 with a slope of zero.

##### Adding set lines

The algorithm to add topologically correct set lines to a grid of circles representing regions runs in three steps.

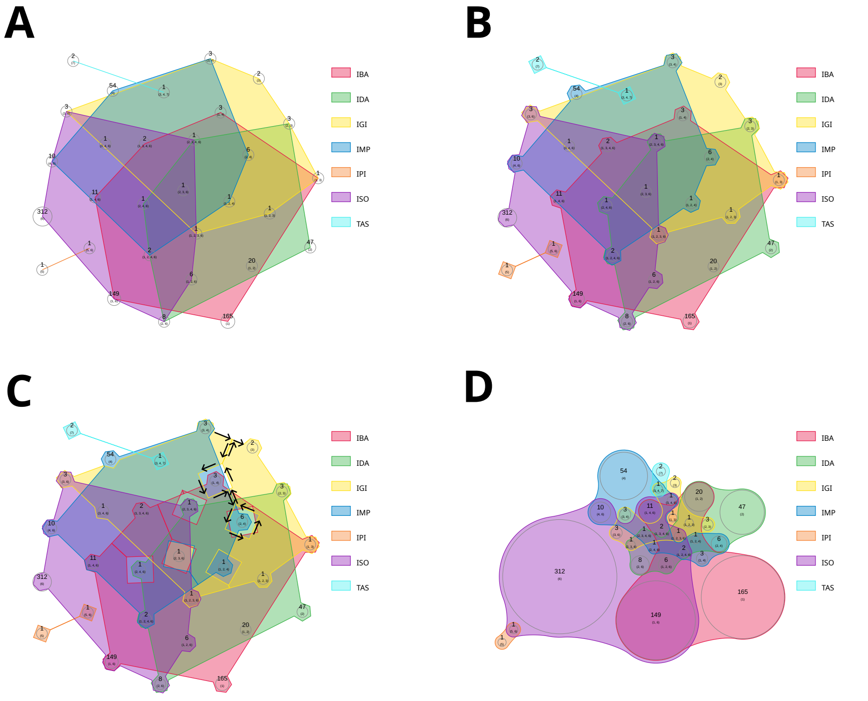


**Figure S1**: Addition of set lines. **A**, addition of set lines that enclose all the regions in each set, and possibly regions not belonging to the set. **B**, addition of lines ensuring that the center of the circles defining the lines are inside the lines. **C**, correction of lines to exclude regions not belonging to each set. Arrows show one example where two regions are excluded from set 3 (IGI, yellow). **D**, final diagram.

##### Adding outer lines

For each set, the procedure iterates over all its regions and selects the leftmost circle (lowest x coordinate, ties do not matter). A reference tangent is created with quadrant 1 and slope 0.

Then, the tangents from the center of this reference circle to the center of every other circle in the set are calculated, and the reference tangent is subtracted from each of them. The circle with the largest resulting tangent is then selected, and a segment is added from the center of the reference circle to the center of the selected circle. Then, the resulting tangent replaces the reference tangent, and the selected circle replaces the reference circle. This procedure is repeated until the new selected circle is the first reference circle and the set line is closed (**Figure S1A**).

##### Including centers

To ensure that the centers of each circle sit inside their sets, the intersections between the segments added in the previous step and the reference circles are calculated. Then, three segments that connect those intersections through the outside of the circle are added (**Figure S1B**).

##### Fixing topology

The lines added in the previous step contain every circle in each set, but they may also contain circles that do not belong to that set. For each circle that does not belong to a set, the distance from its center to each of the set segments is calculated. If the closest point is not contained in the segment, the segment is discarded. The closest segment is selected, and split along the closest point. Six segments are added to the split to connect it again through the outside of the circle, which, after simulation, yields smooth set lines (**Figure S1C-D**).

#### Steps

##### 1 - Attract

Let the Venn diagram contain $n_s$ sets. Each region in this diagram has a binary representation $b_i$ with $n_s$ bits expressing the sets it belongs and does not belong to. Each non-empty region is represented by a circle $c_i$ with a radius $r_i$ such that the area of the circle is proportional to the number of elements in this region. For each $c_i$ and $c_j$ circle pairs, relationships $f_{i,j}$ are calculated as the number of bit positions where both $b_i$ and $b_j$ contain a value of 1.

Briefly, circles representing regions are pseudorandomly placed on a lattice. Then a simulation is executed where circles are subjected to forces depending on their relationships and distance. For two circles $c_i$ and $c_j$ with a relationship of $f_{i,j}$ and radii $r_i$ and $r_j$ at a distance of $d_{i,j}$,

$$F_{i,j} = -k{f_{i,j}}^2 (d_{i,j} - r_i - r_j) + R$$

The constant parameters $k$ and $R$ have been chosen to minimize the number of simulation cycles needed to finish this step. When updating the position of each circle $c_i$, a second friction force $F_f = -k_2 v_i$, is added, where $v_i$ is the speed of $c_i$ and $k_2$ is a constant parameter.

The simulation is run until $\max(v_i)$ is lower than a pre-defined value.

##### 2 - Disperse

This step is very similar to step 1, but parameters $f_{i,j}$ are set to 1 for every pair and the value of $R$ is much higher. As a result, all the forces are repulsive. The simulation is run until $\min(d_i) > \max(r_i)$.

##### 3 - Minimize Compactness

Compactness ($C$) is defined as:

$$C = \sum_{i=0}^{n_r-1} \sum_{j=i+1}^{n_r} (\ln(f_{i,j}) + 2\ln(d_{i,j}))$$

Here, $n_r$ is the number of non-empty regions in the Venn diagram. After calculating the first reference value of $C$, the algorithm iterates over each circle pair. At each iteration, the positions of both circles are swapped and $C$ is calculated again. If the value after the exchange is lower than before, the swap is conserved and the reference is set to the new $C$ value. If not, the swap is undone. The procedure is repeated indefinitely, cycling over the pairs multiple times if necessary, until the reference value has not changed for $n_r$ consecutive iterations.

##### 4 - Minimize Crossings

The procedure is the same as in step 3, but the number of line crossings ($X$) is used instead of compactness. To count crossings, each segment $p_1 \rightarrow p_2$ of a set line is compared to every segment $p_3 \rightarrow p_4$ from every other set line. First, a fast bounding box check filters segments that trivially do not cross. Then, for those comparisons where the bounding boxes of the segments overlap, four tangent objects are created:

$$\begin{aligned} t_1 &= tangent(p_1, p_3) \\ t_2 &= tangent(p_3, p_2) \\ t_3 &= tangent(p_2, p_4) \\ t_4 &= tangent(p_4, p_1) \end{aligned}$$

Both segments are considered as crossing if $((t_2 \leq t_3) \wedge (t_3 \leq t_4))$ or if $((t_3 \leq t_2) \wedge (t_4 \leq t_3))$. With this strategy, it is relatively simple to deal with special cases, such as two points with identical coordinates or segments that intersect at one of the segment points.

##### 5 - Contract Lines

A simulation is built where each segment in each set line becomes a damped spring which sets a force for both segment points with magnitude $F = -kd + bv$, where $k$ and $b$ are constants, $d$ is the distance between the points and $v$ is the speed of the point. Contacts between the segments and the circles representing regions are also calculated.

In this setting, the diagram gets compacted as set lines shrink. Since the simulation depends on a time span in which motions are assumed linear, there are situations in which contacts will not be computed, as during that time span a segment may completely pass through a circle. For this reason, every 50 simulation cycles the topology of the diagram is assessed.

To calculate whether a circle is inside a set line, a horizontal segment is considered from the center of the circle to a point outside the bounding box of the set. This segment is compared to each segment of the set line to find crossings. If the number of segment crossings is odd, the circle is inside the set line. If a circle occupies an incorrect space, the simulation is returned to a previous topologically correct state, the time span is halved and the simulation continues. The correct state may be the initial state if necessary. If the topology of the diagram is correct, the current state substitutes the previous topologically correct state.

The step is finished when the 100-cycle average compactness is higher than the previous value and the compactness does not improve during 50 cycles. The state with the best compactness is saved and used for the following step.

##### 6 - Compact regions

This step is very similar to the previous one, the difference being that a force between circles decaying with the inverse of the distance is added. This force further compacts the circles. The step is finished under the same conditions as step 5.

##### 7 - Smooth set lines

For this step, circles are kept fixed at the positions at the end of step 6. The number of segments in set lines is increased by interpolation and a simulation is set with a damped spring-like force between segment points and an inverse-distance force between segment points and circles. This wraps lines more tightly around circles and makes the areas in the sets more similar to the areas in the regions.

The step is finished when the sum of the areas of the sets has not decreased for 50 cycles.
